## Supplemental Tables for "N-acetylation of α-synuclein enhances synaptic vesicle clustering mediated by α-synuclein and lysophosphatidylcholine"

**Table S1** Identified cross-linked peptides of sample 1 between $^{15}$N-Ac-α-syn and $^{14}$N-Ac-α-syn in LPC (protein: lipid= 1: 50, mol: mol).

| Score | Δ (ppm) | Res of Pr 1* | Sequence of Pr 1* | Res of Pr2* | Sequence of Pr 2* | Mass (Da) |
|---|---|---|---|---|---|---|
| 1.43E-20 | -0.67 | 12 | A$^{11}\underline{K}$EGVVAAAEK$^{21}$ | 23 | T$^{22}\underline{K}$QGVAEAAGK$^{32}$ | 2268.2324 |
| 5.94E-14 | 0.65 | 32 | Q$^{24}$GVAEAAG$\underline{K}$TK$^{34}$ | 10 | G$^{9}$LS$\underline{K}$AK$^{12}$ | 1799.0151 |
| 8.04E-13 | -0.04 | 34 | T$^{33}\underline{K}$EGVLYVGSK$^{43}$ | 23 | T$^{22}\underline{K}$QGVAEAAGK$^{32}$ | 2376.2899 |
| 2.17E-12 | -1.77 | 32 | T$^{22}$KQGVAEAAG$\underline{K}$TK$^{34}$ | 23 | T$^{22}\underline{K}$QGVAEAAGK$^{32}$ | 2484.3546 |
| 5.89E-12 | 0.46 | 45 | T$^{44}\underline{K}$EGVVHGVATVAEK$^{58}$ | 12 | T$^{22}\underline{K}$QGVAEAAGK$^{32}$ | 2733.4911 |
| 1.33E-11 | 0.22 | 12 | A$^{11}\underline{K}$EGVVAAAEK$^{21}$ | 97 | $\underline{K}^{97}$DQLGK$^{102}$ | 1897.0519 |
| 2.22E-11 | -2.07 | 58 | T$^{44}$KEGVVHGVATVAE$\underline{K}$TK$^{60}$ | 10 | G$^{9}$LS$\underline{K}$AK$^{12}$ | 2493.4165 |
| 1.49E-10 | -2.77 | 23 | T$^{22}\underline{K}$QGVAEAAGK$^{32}$ | 97 | $\underline{K}^{97}$DQLGK$^{102}$ | 1884.0315 |
| 2.94E-10 | -1.25 | 34 | T$^{33}\underline{K}$EGVLYVGSK$^{43}$ | 12 | A$^{11}\underline{K}$EGVVAAAEK$^{21}$ | 2389.3103 |
| 1.42E-08 | -1.10 | 21 | E$^{13}$GVVAAAE$\underline{K}$TK$^{23}$ | 10 | G$^{9}$LS$\underline{K}$AK$^{12}$ | 1842.0461 |
| 2.07E-08 | -0.67 | 58 | E$^{46}$GVVHGVATVAE$\underline{K}$TKEQVTNVGGAVVTGVTAVAQK$^{80}$ | 12 | G$^{7}$LSKA$\underline{K}$EGVVAAAEK$^{21}$ | 5027.7505 |
| 3.43E-08 | -2.37 | 43 | T$^{33}$KEGVLYVGS$\underline{K}$TK$^{45}$ | 10 | G$^{9}$LS$\underline{K}$AK$^{12}$ | 2149.2357 |
| 5.39E-07 | 2.62 | 43 | T$^{33}$KEGVLYVGS$\underline{K}$TK$^{45}$ | 43 | T$^{33}$KEGVLYVGS$\underline{K}$TK$^{45}$ | 2955.6531 |

Δ: mass error for crosslink assignments; Res: crosslinked residue (sequence number) within corresponding proteins; Pr: protein; Underlined K in sequence is the cross-linked lysine.

*The crosslinked pair of residues were from two individual Ac-α-syn: $^{14}$N-Ac-α-syn and $^{14}$N-Ac-α-syn, $^{14}$N-Ac-α-syn and $^{15}$N-Ac-α-syn, $^{15}$N-Ac-α-syn and $^{14}$N-Ac-α-syn, or $^{15}$N-Ac-α-syn and $^{15}$N-Ac-α-syn. Detailed data processing procedure could be referred to Methods session.

**Table S2** Identified cross-linked peptides of sample 2 between $^{15}$N-Ac-α-syn and $^{14}$N-Ac-α-syn in LPC (protein: lipid= 1: 50, mol: mol).

| Score | Δ (ppm) | Res of Pr 1* | Sequence of Pr 1* | Res of Pr2* | Sequence of Pr 2* | Mass (Da) |
|---|---|---|---|---|---|---|
| 2.56E-17 | -0.42 | 45 | T$^{44}\underline{\text{K}}$EGVVHGVATVAEK$^{58}$ | 12 | A$^{11}\underline{\text{K}}$EGVVAAAEK$^{21}$ | 2733.4911 |
| 3.92E-15 | 0.45 | 12 | A$^{11}\underline{\text{K}}$EGVVAAAEK$^{21}$ | 23 | T$^{22}\underline{\text{K}}$QGVAEAAGK$^{32}$ | 2268.2324 |
| 3.41E-14 | 0.97 | 34 | T$^{33}\underline{\text{K}}$EGVLYVGSK$^{43}$ | 23 | T$^{22}\underline{\text{K}}$QGVAEAAGK$^{32}$ | 2376.2899 |
| 2.82E-13 | 0.45 | 32 | Q$^{24}$GVAEAAG$\underline{\text{K}}$TK$^{34}$ | 10 | G$^{7}$LS$\underline{\text{K}}$AK$^{10}$ | 1799.0151 |
| 6.88E-13 | 0.59 | 43 | T$^{33}$KEGVLYVGS$\underline{\text{K}}$TK$^{45}$ | 12 | A$^{11}\underline{\text{K}}$EGVVAAAEK$^{21}$ | 2618.4529 |
| 4.51E-12 | 1.04 | 12 | A$^{11}\underline{\text{K}}$EGVVAAAEK$^{21}$ | 97 | $\underline{\text{K}^{97}}$DQLGK$^{102}$ | 1897.0519 |
| 2.59E-11 | -2.90 | 23 | T$^{22}\underline{\text{K}}$QGVAEAAGK$^{32}$ | 97 | $\underline{\text{K}^{97}}$DQLGK$^{102}$ | 1884.0315 |
| 1.28E-10 | 0.36 | 32 | Q$^{24}$GVAEAAG$\underline{\text{K}}$TK$^{34}$ | 97 | $\underline{\text{K}^{97}}$DQLGK$^{102}$ | 1884.0315 |
| 1.07E-09 | -1.42 | 21 | A$^{11}$KEGVVAAAE$\underline{\text{K}}$TK$^{23}$ | 12 | A$^{11}\underline{\text{K}}$EGVVAAAEK$^{21}$ | 1965.9903 |
| 2.81E-09 | 2.82 | 21 | E$^{13}$GVVAAAE$\underline{\text{K}}$TK$^{23}$ | 10 | G$^{7}$LS$\underline{\text{K}}$AK$^{10}$ | 2510.3954 |
| 7.27E-08 | 0.30 | 43 | T$^{33}$KEGVLYVGS$\underline{\text{K}}$TK$^{45}$ | 97 | $\underline{\text{K}^{97}}$DQLGK$^{102}$ | 1842.0461 |

Δ: mass error for crosslink assignments; Res: crosslinked residue (sequence number) within corresponding proteins; Pr: protein; Underlined K in sequence is the cross-linked lysine.

*The crosslinked pair of residues were from two individual Ac-α-syn: $^{14}$N-Ac-α-syn and $^{14}$N-Ac-α-syn, $^{14}$N-Ac-α-syn and $^{15}$N-Ac-α-syn, $^{15}$N-Ac-α-syn and $^{14}$N-Ac-α-syn, or $^{15}$N-Ac-α-syn and $^{15}$N-Ac-α-syn. Detailed data processing procedure could be referred to Methods session.

**Table S3** Identified cross-linked peptides of sample 3 between $^{15}$N-Ac-α-syn and $^{14}$N-Ac-α-syn in LPC (protein: lipid= 1: 50, mol: mol).

| Score | Δ (ppm) | Res of Pr 1* | Sequence of Pr 1* | Res of Pr2* | Sequence of Pr 2* | Mass (Da) |
|---|---|---|---|---|---|---|
| 3.49E-15 | 0.21 | 12 | A$^{11}$KEGVVAAAEK$^{21}$ | 23 | T$^{22}$KQGVAEAAGK$^{32}$ | 2268.2324 |
| 1.32E-13 | 3.78 | 60 | T$^{59}$KEQVTNVGGAVVTGVTAVAQK$^{80}$ | 34 | T$^{33}$KEGVLYVGSK$^{43}$ | 3473.8980 |
| 4.26E-13 | 2.34 | 34 | T$^{33}$KEGVLYVGSK$^{43}$ | 23 | T$^{22}$KQGVAEAAGK$^{32}$ | 2376.2899 |
| 1.14E-12 | 0.67 | 12 | A$^{11}$KEGVVAAAEK$^{21}$ | 97 | K$^{97}$DQLGK$^{102}$ | 1897.0519 |
| 2.32E-12 | 1.24 | 45 | T$^{44}$KEGVVHGVATVAEK$^{58}$ | 23 | T$^{22}$KQGVAEAAGK$^{32}$ | 2720.4707 |
| 5.94E-12 | 1.20 | 96 | T$^{81}$VEGAGSIAAATGFVKK$^{97}$ | 12 | A$^{11}$KEGVVAAAEK$^{21}$ | 2815.5330 |
| 1.07E-10 | 0.52 | 43 | T$^{33}$KEGVLYVGSKTK$^{45}$ | 12 | A$^{11}$KEGVVAAAEK$^{21}$ | 2618.4529 |
| 5.04E-09 | 0.91 | 21 | A$^{11}$KEGVVAAAEKTK$^{23}$ | 97 | K$^{97}$DQLGK$^{102}$ | 2126.1946 |
| 2.29E-08 | 1.40 | 96 | T$^{91}$VEGAGSIAAATGFVKK$^{97}$ | 23 | T$^{22}$KQGVAEAAGK$^{32}$ | 2802.5126 |
| 1.52E-07 | 1.15 | 43 | T$^{33}$KEGVLYVGSKTK$^{45}$ | 10 | G$^{7}$LSKAK$^{10}$ | 2149.2357 |
| 2.52E-07 | -3.62 | 34 | T$^{33}$KEGVLYVGSK$^{43}$ | 12 | A$^{11}$KEGVVAAAEK$^{21}$ | 2389.3103 |
| 5.32E-07 | -2.34 | 60 | T$^{59}$KEQVTNVGGAVVTGVTAVAQK$^{80}$ | 21 | E$^{13}$GVVAAAEKTK$^{23}$ | 3395.8510 |
| 7.21E-07 | 0.58 | 12 | A$^{11}$KEGVVAAAEK$^{21}$ | 12 | A$^{11}$KEGVVAAAEK$^{21}$ | 2281.2528 |
| 7.33E-07 | 2.58 | 32 | T$^{22}$KQGVAEAAGKTK$^{34}$ | 23 | T$^{22}$KQGVAEAAGK$^{32}$ | 2484.3546 |
| 8.89E-07 | 0.99 | 21 | E$^{13}$GVVAAAEKTK$^{23}$ | 10 | G$^{7}$LSKAK$^{10}$ | 1842.0461 |

Δ: mass error for crosslink assignments; Res: crosslinked residue (sequence number) within corresponding proteins; Pr: protein; Underlined K in sequence is the cross-linked lysine.

*The crosslinked pair of residues were from two individual Ac-α-syn: $^{14}$N-Ac-α-syn and $^{14}$N-Ac-α-syn, $^{14}$N-Ac-α-syn and $^{15}$N-Ac-α-syn, $^{15}$N-Ac-α-syn and $^{14}$N-Ac-α-syn, or $^{15}$N-Ac-α-syn and $^{15}$N-Ac-α-syn. Detailed data processing procedure could be referred to Methods session.

**Table S4** Identified cross-linked peptides of sample 1 between $^{15}$N-Ac-α-syn and $^{14}$N-Ac-α-syn in DOPS (protein: lipid= 1: 50, mol: mol).

| Score | Δ (ppm) | Res of Pr 1* | Sequence of Pr 1* | Res of Pr2* | Sequence of Pr 2* | Mass (Da) |
|---|---|---|---|---|---|---|
| 2.08E-16 | 0.45 | 32 | Q$^{24}$GVAEAAGKTK$^{34}$ | 10 | G$^{7}$LSKAK$^{10}$ | 1799.0151 |
| 2.67E-14 | 2.65 | 45 | T$^{44}$KEGVVHGVATVAEK$^{58}$ | 23 | T$^{22}$KQGVAEAAGK$^{32}$ | 2720.4707 |
| 4.22E-14 | -2.67 | 45 | T$^{44}$KEGVVHGVATVAEK$^{58}$ | 12 | A$^{11}$KEGVVAAAEK$^{21}$ | 2733.4911 |
| 1.76E-12 | -0.66 | 12 | A$^{11}$KEGVVAAAEK$^{21}$ | 23 | T$^{22}$KQGVAEAAGK$^{32}$ | 2268.2324 |
| 1.92E-09 | -0.87 | 21 | E$^{13}$GVVAAAEKTK$^{23}$ | 23 | T$^{22}$KQGVAEAAGK$^{32}$ | 2298.2430 |
| 8.17E-09 | 1.29 | 21 | A$^{11}$KEGVVAAAEKTK$^{23}$ | 10 | G$^{7}$LSKAK$^{10}$ | 2041.1782 |

Δ: mass error for crosslink assignments; Res: crosslinked residue (sequence number) within corresponding proteins; Pr: protein; Underlined K in sequence is the cross-linked lysine.

*The crosslinked pair of residues were from two individual Ac-α-syn: $^{14}$N-Ac-α-syn and $^{14}$N-Ac-α-syn, $^{14}$N-Ac-α-syn and $^{15}$N-Ac-α-syn, $^{15}$N-Ac-α-syn and $^{14}$N-Ac-α-syn, or $^{15}$N-Ac-α-syn and $^{15}$N-Ac-α-syn. Detailed data processing procedure could be referred to Methods session.

**Table S5** Identified cross-linked peptides of sample 2 between $^{15}$N-Ac-α-syn and $^{14}$N-Ac-α-syn in DOPS (protein: lipid= 1: 50, mol: mol).

| **Score** | **Δ (ppm)** | **Res of Pr 1*** | **Sequence of Pr 1*** | **Res of Pr2*** | **Sequence of Pr 2*** | **Mass (Da)** |
|---|---|---|---|---|---|---|
| 3.01E-15 | -0.73 | 43 | T$^{33}$KEGVLYVGS$\underline{K}$TK$^{45}$ | 43 | E$^{35}$GVLYVGS$\underline{K}$TK$^{45}$ | 2726.5105 |
| 1.71E-11 | 2.06 | 96 | T$^{81}$VEGAGSIAAATGFV$\underline{K}$K$^{97}$ | 12 | A$^{11}$$\underline{K}$EGVVAAAEK$^{21}$ | 2815.5330 |
| 1.22E-10 | -2.90 | 32 | Q$^{24}$GVAEAAG$\underline{K}$TK$^{34}$ | 97 | $\underline{K^{97}}$DQLGK$^{102}$ | 1884.0315 |
| 3.80E-08 | -0.35 | 96 | T$^{81}$VEGAGSIAAATGFV$\underline{K}$K$^{97}$ | 10 | G$^{7}$LS$\underline{K}$AK$^{10}$ | 2346.3157 |
| 4.64E-07 | 0.38 | 23 | T$^{22}$$\underline{K}$QGVAEAAGK$^{32}$ | 23 | T$^{22}$$\underline{K}$QGVAEAAGK$^{32}$ | 2255.2120 |

Δ: mass error for crosslink assignments; Res: crosslinked residue (sequence number) within corresponding proteins; Pr: protein; Underlined K in sequence is the cross-linked lysine.

*The crosslinked pair of residues were from two individual Ac-α-syn: $^{14}$N-Ac-α-syn and $^{14}$N-Ac-α-syn, $^{14}$N-Ac-α-syn and $^{15}$N-Ac-α-syn, $^{15}$N-Ac-α-syn and $^{14}$N-Ac-α-syn, or $^{15}$N-Ac-α-syn and $^{15}$N-Ac-α-syn. Detailed data processing procedure could be referred to Methods session.

**Table S6** Identified cross-linked peptides of sample 3 between $^{15}$N-Ac-α-syn and $^{14}$N-Ac-α-syn in DOPS (protein: lipid= 1: 50, mol: mol).

| Score | Δ (ppm) | Res of Pr 1* | Sequence of Pr 1* | Res of Pr2* | Sequence of Pr 2* | Mass (Da) |
|---|---|---|---|---|---|---|
| 4.55E-14 | 0.36 | 32 | Q$^{24}$GVAEAAG$\underline{K}$TKEGVLYVGSK$^{43}$ | 21 | A$^{11}$KEGVVAAAE$\underline{K}$TK$^{23}$ | 3429.8717 |
| 9.60E-14 | 0.25 | 32 | Q$^{24}$GVAEAAG$\underline{K}$TK$^{34}$ | 10 | G$^{7}$LS$\underline{K}$AK$^{10}$ | 1799.0151 |
| 1.05E-07 | -1.97 | 45 | T$^{44}\underline{K}$EGVVHGVATVAEK$^{58}$ | 43 | E$^{35}$GVLYVGS$\underline{K}$TK$^{45}$ | 2841.5486 |
| 1.08E-07 | 1.16 | 23 | T$^{22}\underline{K}$QGVAEAAGK$^{32}$ | 32 | Q$^{24}$GVAEAAG$\underline{K}$TK$^{34}$ | 2255.2120 |
| 2.30E-07 | -1.49 | 96 | T$^{81}$VEGAGSIAAATGFV$\underline{K}$K$^{97}$ | 12 | A$^{11}\underline{K}$EGVVAAAEK$^{21}$ | 2815.5330 |
| 5.89E-07 | 3.15 | 6 | M$^{1}$DVFM$\underline{K}$GLSK$^{10}$ | 32 | Q$^{24}$GVAEAAG$\underline{K}$TK$^{34}$ | 2351.2228 |

Δ: mass error for crosslink assignments; Res: crosslinked residue (sequence number) within corresponding proteins; Pr: protein; Underlined K in sequence is the cross-linked lysine.

*The crosslinked pair of residues were from two individual Ac-α-syn: $^{14}$N-Ac-α-syn and $^{14}$N-Ac-α-syn, $^{14}$N-Ac-α-syn and $^{15}$N-Ac-α-syn, $^{15}$N-Ac-α-syn and $^{14}$N-Ac-α-syn, or $^{15}$N-Ac-α-syn and $^{15}$N-Ac-α-syn. Detailed data processing procedure could be referred to Methods session.
